# On the mechanism of long-range orientational order of fibroblasts

**DOI:** 10.1101/119669

**Authors:** Xuefei Li, Rajesh Balagam, Ting-Fang He, Peter P. Lee, Oleg A. Igoshin, Herbert Levine

**Author notes:** X.L., R.B., O.A.I., and H.L. designed research; X.L., R.B. and T.H. performed research; and all authors wrote the paper. We declare no conflict of interest. X.L. and R.B. contributed equally to this work.

## Abstract

Long-range alignment ordering of fibroblasts have been observed in the vicinity of cancerous tumors and can be recapitulated with *in vitro* experiments. However, the mechanisms driving their ordering are not understood. Here we show that local collision-driven nematic alignment interactions among fibroblasts are insufficient to explain observed long-range alignment. One possibility is that there exists another orientation field co-evolving with the cells and reinforcing their alignment. We propose that this field reflects the mechanical cross-talk between the fibroblasts and the underlying fibrous material on which they move. We demonstrate that this new long-range interaction can give rise to high nematic order and to the observed patterning of the cancer microenvironment.

**Significance Statement:** Long-range alignment patterns of fibroblasts have been observed both *in vivo* and *in vitro*. However, there has not been much understanding of the underlying mechanism. In this work, we demonstrate that these patterns cannot be simply explained by their steric interaction with one another during collisions. Instead, we propose that fibroblasts may collectively align through non-local interactions arising from their modification of an underlying extracellular matrix. The proposed mechanism explains the observed co-alignment between fibroblasts and collagen fibers around tumors and can be be tested in future experiments that can image the dynamics of this pattern formation *in vivo* or *in vitro*

---

Fibroblasts are spindle shaped cells that are highly motile and are involved in many critical biological processes such as wound healing (1, 2). Recently, their major role in shaping the local micro-environment around a growing tumor was shown in numerous studies (3–5). As a result, fibroblasts can affect the ability of cancer cells to metastasize (6–8) and conversely the ability of the immune system to find and attack those cells (9).

An isolated fibroblast moves back and forth on cover slips for over 60 hours without significant change of the direction of its major axis (10). Typically, those fibroblasts are in a spindle shape with an aspect ratio from 2 to 5. Apart from steric constraints, fibroblasts barely interact with each other (10). Curiously, imaging of tumor micro-environments often indicates long-range ordering of fibroblasts. This order often takes the form of circumferential alignment of the cells in a region surrounding the cancer cells (11, 12). The mechanisms that lead to this ordering are not well understood.

Notably, a similar ordering can be observed for underlying collagen fibers (6, 13). Collagen fibers are the main structural component in the extracellular space of various normal connective tissues and play a significant role in local cancer cell invasion and in metastasis (14), Aligned fibers perpendicular to the boundary cancer-cell clusters facilitate local invasions of cancer cells, and conversely aligned fibers parallel to the tumor boundary may restrict the migration of cancer cells. Furthermore, the circumferential collagen fiber structure has been hypothesized to be responsible for the observed separation between immune and cancer cells (13). Specifically, *ex vivo* assays indicate that the migration of tumor-killing CD8^+^T cells is reduced where dense collagen fibers form conduit-like structures (15). Therefore, understanding the mechanism of the pattern formation of fibroblasts and collagen fibers in the cancer microenvironment is important to strike a balance between constraining cancer cell from invasion and enabling the infiltration of cancer-killing immune cells.

*In vivo* experimental observation of this pattern formation dynamically is technically challenging. On the other hand, long-range ordering of fibroblasts can be recapitulated in two dimensional cell culture experiments. In particular, Duclos et al. (10) generated quantitative measurements of cluster orientation as fibroblasts move, collide, and proliferate (grow and divide) on cover slips in 2D. To quantify the collective alignment of fibroblast, the authors used their data to compute the population averaged nematic order parameter 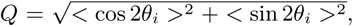 which increases from 0 for randomly oriented cell population up to the value of 1 for perfectly aligned cells. The results indicate that as cell density increases due to cell growth the cells become more aligned and the order parameter increases. After roughly 80 hrs, cell motions become significantly limited due to jamming and the nematic order parameter freezes at a fixed value. Fibroblasts grown in a relatively narrow channels (100 cells across), robustly display almost perfect nematic alignment (order parameter *Q* ≈ 1).

In this work we aim to understand the underlying mechanisms of the long-range nematic alignment of fibroblasts. First, noting that steric interactions between fibroblast may lead to local alignment, we ask if these interactions are sufficient to explain the experimental results described above. The results indicate that perfect long-range alignment can not emerge from models treating fibroblasts as apolar active particles with hard collision interactions (active nematics). Therefore, we further test models in which non-local interactions between fibroblasts are introduced. Biologically, these models are motivated by the observations (16–19) that fibroblasts are able to deposit and reorganize fibril materials around them. Using this model, we investigated whether coupling between cells and their mechanical environment can recapitulate patterns of fibroblasts both *in vitro* and *in vivo*.

## Results

### “Fibroblasts” interacting through collisions can not produce the long-range alignment

We first set out to test whether the steric interactions between fibroblasts are sufficient to explain the long-range alignment of cells observed in the experiment. To proceed, we start with a Monte Carlo simulation framework, in which fibroblasts are represented as hard elliptic particles. Particles move back and forth and re-orientate after overlapping with neighbors, Fig. 1A. To mimic the fibroblast proliferation observed in experiments (10), new particles are added in front of randomly selected particles and with the same orientation as the selected particle. In addition, we also studied another model that considers explicit forces acting on particles. In this model, each fibroblast cell is represented by a 2-dimensional sphero-cylinder, Fig. 1B. Implementation details of various cell processes are described in *Material and Methods*.

**Fig. 1.**
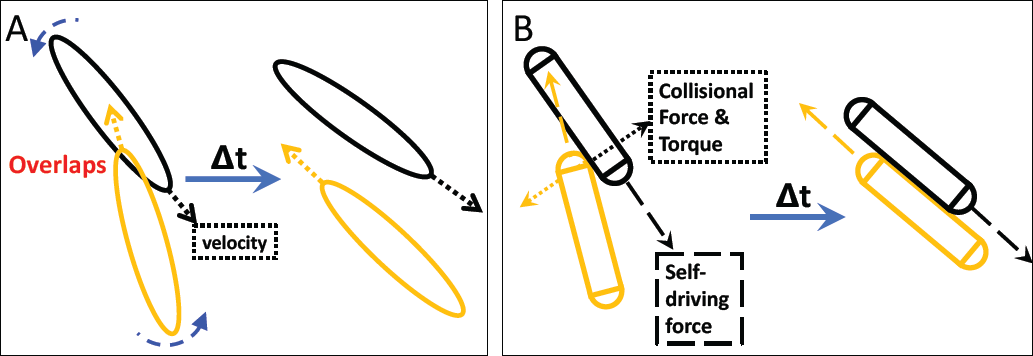
Illustration of the modeling approaches used in this study. Both simulation frameworks describe the effective steric interactions and can result in the local alignment of “fibroblasts”. (A) In the Monte Carlo simulation framework, fibroblasts are treated as hard elliptic particles. Each “fibroblast” moves along its long (major) axis with a constant velocity and can reverse its direction. If fibroblasts overlap with neighbors, they rotate by a randomly selected angle within a fixed range. (B) In the Newtonian dynamic simulation framework, a “fibroblast” is represented by a sphero-cylinder. It moves along its long axis driven by a constant self-driving force whose direction can also reverse. A constant collisional force is applied after each contact and the consequent torques drive the rotations of “fibroblasts”. Detailed descriptions of both simulations are presented in the *Materials and Methods* section.

With the collision-like interaction, our simulations (Fig. 2) did not reproduce the perfect alignment of fibroblasts (particles in the model) in a channel-like structure, where the width of the channel is 60 fibroblasts across. In our simulations, while those particles right next to channel boundaries align parallel to boundaries, this alignment disappears for cells in the channel center, Fig. 2A and 2B. The spatial-dependent alignment is further quantified in Fig. S18. In contrast, experiments in (10) showed nearly-perfect alignment of fibroblasts to the channel boundaries up-to a channel width of 500*µm* (around 100 cells across), where the width of a fibroblast is around 5*µm*, Fig. 3B. We note, that while changes in the particle geometry, e.g. aspect ratio (ratio between length and width of a particle), can affect the alignment, perfect nematic alignment is never observed for biologically relevant parameter values, Fig. 2C.

**Fig. 2.**
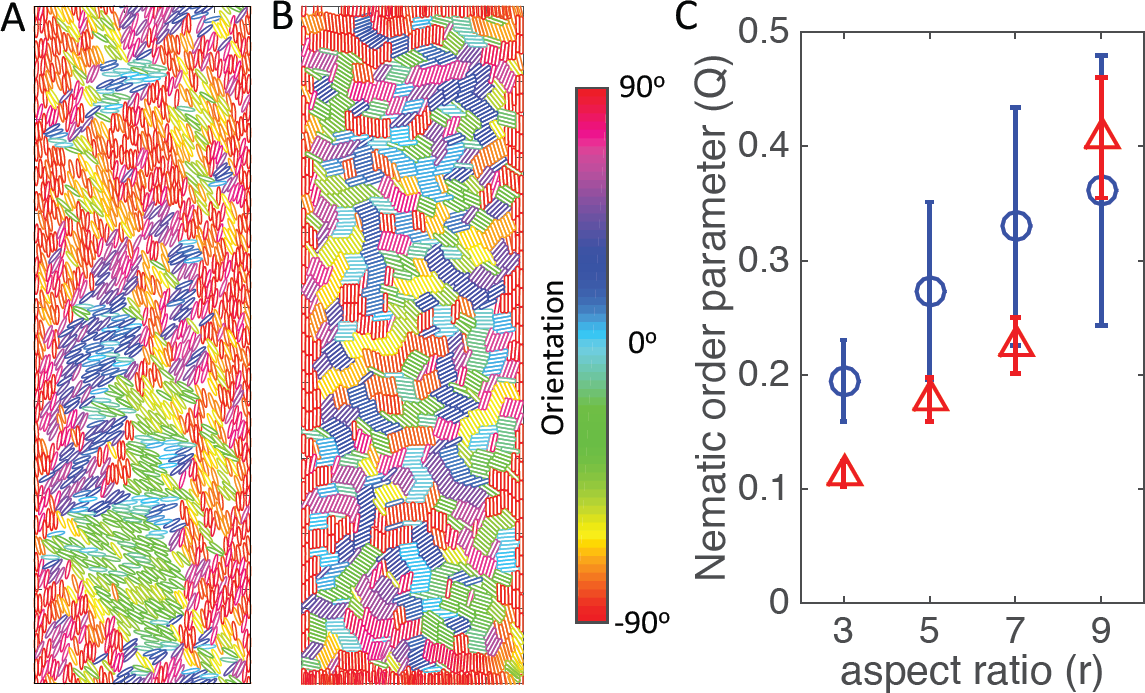
(A) and (B) Snapshots of the simulation at step 2000 for ellipses (A) or sphero-cylinders (B). Color code for the orientation of particles is displayed. (C) Nematic order parameter *Q* in a channel. Values of nematic order parameter are shown as circles for elliptic particles or as triangles for sphero-cylinders. In these simulations, channels are set as 60 cells across horizontal axis. Periodic boundary conditions are applied along the vertical axis. Two hard walls are implemented on the left and right boundary. The parameters and detailed simulation procedures can be found in the *Materials and Methods*.

**Fig. 3.**
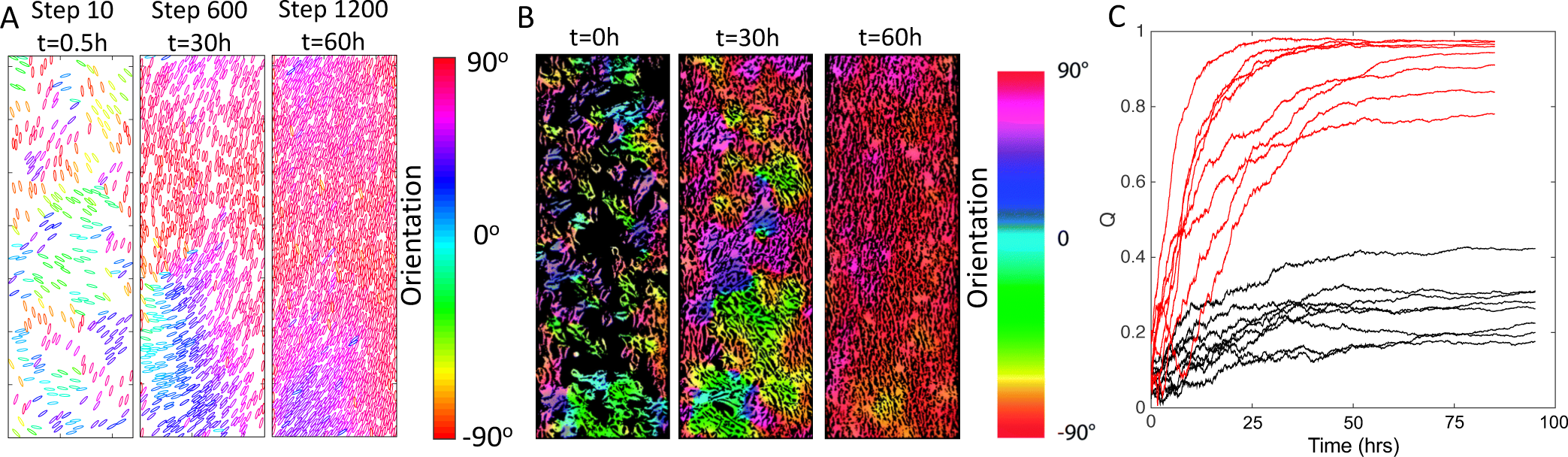
Numerical simulations generating long-range alignment order in a channel-like geometry. The number of particles gradually increases from 300 (packing fraction 0.12) to over 1500 (packing fraction 0.58) in the simulations. The aspect ratio of particles in these examples is 5. (see Materials and Methods) is chosen as 0.1 in those simulations. (A) Snapshots of the particle configuration of one simulation at step 10, 600 and 1200. (B) Evolution of perfect nematic alignment of fibroblasts in a 2D channel *in vitro*. Width of channel is 0.5mm. The figure is adapted from Ref. (10). (C) Different realizations of our Monte Carlo simulation carried out in a channel-like structure. Black lines are different realizations of the system with only collision interactions and the parameters are the same as those in Fig. 2. Red lines represent the evolution of nematic order with multi-alignment interactions between fibroblasts and fibers applied (*R_c_* = 15 *b*, α = 0.1).

We have investigated a number of other modifications of our basic model (see SI). For example, we tested the effects of non-reversal motion, channel width, flexibility of “fibroblasts”, initial cell density, initial alignment orders, etc. However, we did not observe significant change in the results. We therefore conclude that additional mechanisms are needed to explain emergent nematic ordering.

### Mutual alignment between fibroblasts and fibers can generate perfect alignment in narrow channels

Motivated by (20), we propose a novel mechanism in which there exists an orientation field coupled to nearby particles and that this field can in turn affect the orientation of those particles. For fibroblasts, this idea is supported by the fact that cells secrete materials for extracellular matrix (ECM) such as collagen (16, 17). Furthermore, as fibroblasts move, they contract and align ECM fibers depending on their direction of motion (18, 19). Recent experimental results also show that a fiber network under strain over a threshold time can change its structural and properties in a non-reversible fashion (21). As a result, the fiber network at some point in space can “remember” the average orientation of the particles that have previously traversed close to that location (22). We hypothesize that mutual alignment of ECM network and fibroblasts drives the emergence of the patterns in cancer tissue specimens and the high nematic order in culture.

Rather than formulate an extremely detailed mechanical model of ECM-fibroblast co-orientation, we have opted for a simple proof-of-principle phenomenological approach. Due to the qualitative similarity between the Monte Carlo simulation and the explicit-force model, we only implement and study the new mechanism using the Monte Carlo simulation framework. The local orientation field at a point on a 2D surface is affected by the orientations of particles *_n_* within a radius of *R_c_*. However, as the orientation field gradually changes in response to nearby fibroblasts, it maintains a memory of previous motion in its vicinity. On the other hand, each particle gradually turns to align with the direction of the local orientation field and moved according to this new orientation. Detailed implementation of the model can be found in the *Materials and Methods* section.

The model implementing the above-described mechanism, shows higher order of particle alignment. Specifically, nearly-perfect nematic alignment can be realized for the channel-like confinement setup, where the width of the channel is 60 particles across. In this case, cells close to the hard walls move around and orient themselves to align with the wall. Then this orientation propagates into the channel center. The time-dependent configurations of particles are shown in Fig. 3A. These features are quite similar to those reported by Duclos et al. (10), Fig. 3B, and quantified in Fig. 3C. We should note that for the simulations in the channel case, even with mutual-alignment interactions, there are still 2 cases out of 8 simulations which stay at a high but not nearly-perfect orientational order. In those cases, defects developed close to the boundary and did not resolve before the particle density increased to a level that the jamming effect kicked in. As expected, with a bigger channel width, clusters of fibroblast, which do not parallel to the boundary, will develop and the orientational order decreases (Fig. S19).

We also performed simulations with periodic boundary conditions in both x and y directions, which mimics the fibroblast culture experiment on a large 2D surface instead of a channel-like confinement setup. We show that the non-local alignment interaction between “fibroblast” facilitated by the “fiber” field is essential to recapitulate the long-range nematic alignment of fibroblasts on a 2D surface (Fig. S20).

### Mutual alignment between fibroblasts and fibers results in the long-range circumferential alignment of fibroblasts and fibers around obstacles

To investigate whether the mutual alignment model can explain the *in vivo* results, we now add obstacles to our simulation mimicking the effect of the cancer-cell clusters. We found that the ability of these simulations to produce long-range alignment around the obstacles depends on the rules of fibroblast proliferation. If new elliptic particles are introduced into the simulation box with the same probability everywhere, i.e. if fibroblast proliferation rate is not location-dependent, simulations often result in clusters of particles pointing towards an obstacle (see Fig. S21). Since it takes some time before particles align to the boundary of an obstacle and re-organize the orientation field around an obstacle, jamming effects due to high particle density can hinder the circumferential alignment. Therefore, we concluded that it is important to ensure that particles around an obstacle do not jam before they align with the boundary.

To avoid this possible jamming, we introduce one other feature, namely that new elliptic particles be introduced with a higher probability in a range close to the obstacles. This assumption is based again on the observations of (10), that fibroblasts closer to boundaries proliferate faster. This may also be a feature of the cancer micro-environment where fibroblasts can be stimulated by growth factors released by the tumor.

With this new ingredient applied in the simulation, we can now generate the pattern in which the ellipses as well as orientation field align azimuthally around obstacles. These results are shown in Fig. 4A and 4B.

**Fig. 4.**
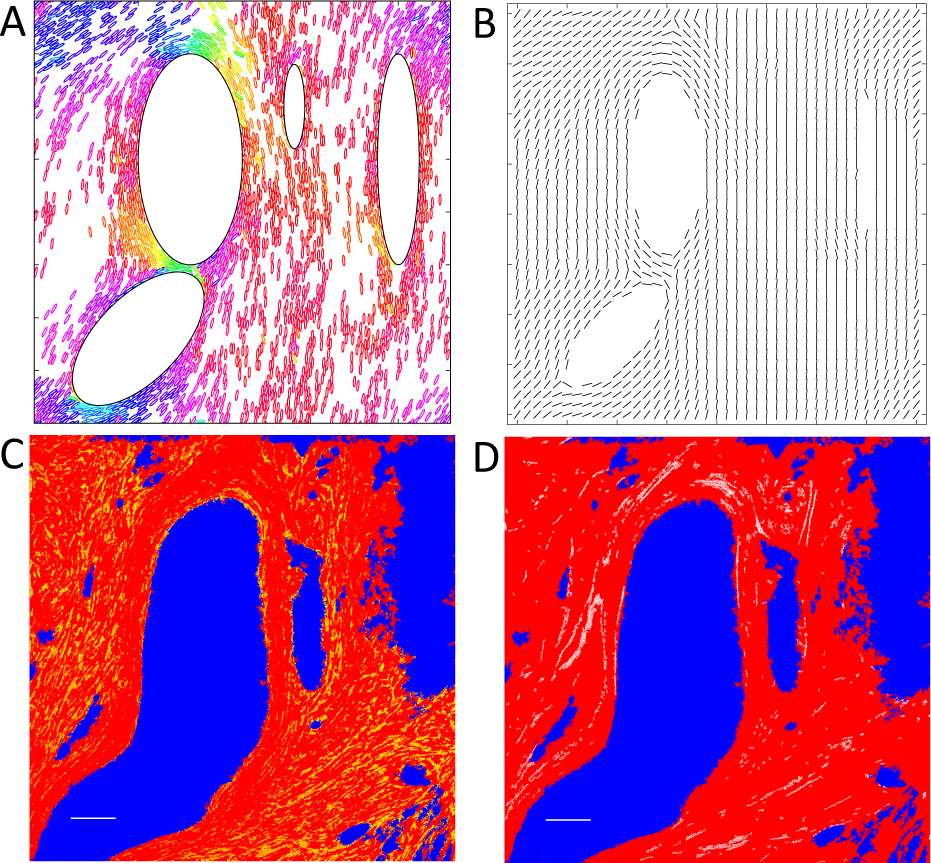
(A) and (B): Numerical simulations with mutual alignment of “fibroblasts” and “fibers”. Hard elliptic obstacles were assigned beforehand. 400 particles were first randomly assigned. New particles are added only within a range close to large obstacles, i.e., half length of a “fibroblast”. Color-coding of elliptic particles are the same as that in Fig. 3A. Configurations of fibroblasts is shown in (A). Orientation field is shown in (B) by the direction of each black line on each grid point. We reduce the number of grid points to make the plot clear. Qualitatively, the simulation results are comparable to experimental result shown in (C) and (D). (C) A breast cancer specimen from a patient. Regions with cancer cells are labeled in blue and background color is set as red for clearer presentation. One particular protein, alpha smooth muscle actin, which is expressed by cancer associated fibroblasts (23) in cancerous tissue, is labeled in yellow. These fibroblasts are found mostly in the space between cancer-cell clusters and orient azimuthally around the clusters. Scale Bar 100 *µm* (white, left corner). (D) The same field of view as in (C). Collagen fibers captured by a Second Harmonic Generation microscope (24) are labeled in white. They surround regions of breast cancer cells clusters (blue) and are parallel to the boundary of the cancer-cell clusters. Scale Bar 100*µm* (white, left corner). Detailed experimental methods on immunohistochemical labeling of the specimens and image acquisition can be found in the supplementary information.

We should note that without the coupling between particles and the orientation field in our simulations, the circumferential pattern of particles can also emerge in the region immediately around the obstacle (Fig. S22). However, as we move away from the obstacle, the circumferential pattern disappears and the particles are not well-organized as shown in the mutual-alignment model (Fig. 4A and Fig. S22).

### Experimental results confirm the predicted circumferential and long-range alignment of fibroblasts and collagen fibers around tumor-cell clusters

When we consider the mutual alignment between fibroblasts and fibers, our model predicts a long-range circumferential alignment of both fibroblasts and collagen fibers around tumor-cell clusters. In order to confirm the prediction in the patient cancer-tissue specimens, we simultaneously imaged cancer-associated fibroblasts and collagen fibers in the same region of patient tumor specimens as shown in Fig. 4C and 4D. The experimentally observed patterns of fibroblasts and fibers closely match the simulation results, which suggest our model can capture the underlying mechanism of the patterns of fibroblasts and fibers in cancer tissue. We should note that the pattern observed in the experiment shown in Fig. 4C and 4D is a projection of the actual 3D structure on a 2D section. Although we have not been able to perform a 3D simulation so far, we argue that the mechanism proposed should be important for the long-range alignment order in 3D cancerous tissues.

## Discussion

In this paper, to understand the underlying mechanism of the long-range alignment pattern of fibroblasts observed both *in vitro* and *in vivo*, we developed computational frameworks to test the roles of steric interactions among neighboring cells and mutual interactions between cells and their ECM network. Our results indicate that mutual alignment between fibroblasts and collagen fibers is critical to recapitulate the observed long-range alignment pattern of fibroblasts.

First, we demonstrated that treating fibroblasts as apolar active particles with hard collision interactions (active nematics) cannot recapitulate the long-range alignment of fibroblasts *in vitro* (Fig. 2). Our results are not totally surprising in light of some previous studies. For point-like particles with nematic interactions described by a Vicsek type model (25), there can be high nematic order in a low rotational noise regime. However, the results are different for hard eliptic particles. Shi and Ma (26) performed kinetic Monte-Carlo simulations of such particles moving back and forth and colliding on the 2D surface, which mimics experiments carried out by Narayan et al. (27). The results indicate that, unlike particles performing Brownian motions, nematic order for active nematics gradually decreases with increase in particle density. Direction reversal of self-propelled particles can also play a key role, as it can disrupt nematic order or cell-clustering (20, 28). There also have been recent analytical work (29, 30) showing that highly ordered state can be unstable in active nematics.(31) We also tested another idea where fibroblasts can secrete matrix metalloproteinases (MMPs), which can degrade fiber network and create a channel for fibroblasts to follow and move inside. However, this mechanism alone is not able to generate the long-range order either. Detailed implementation of this mechanism and results can be found in the supplementary information (Fig. S23).

Furthermore, we propose a new mechanism that explains long-range ordering patterns of fibroblasts: the nearly-perfect nematic alignment in 2D channel cultures (Fig. 3B) and the azimuthal alignment around cancer-cell clusters (Fig. 4C). Specifically, we propose that fibroblasts may align with each other through non-local interactions arising from their modification of an underlying ECM network. We demonstrated that such a long-range interaction is necessary to even qualitatively reproduce experimental observations (e.g., Fig. 3C). We note that previous studies in other contexts have indicated that mutual mechanical interactions between cells and their micro-environment can be very important to understand certain types of cell-pattern formation. For example, the alignment of basement-membrane fibers is driven by the collective rotation of cells in the Drosophila embryo of fruit fly (32).

Our studies serve as an initial attempt to explore the mechanical aspects of collective cell motion and pattern formation. The proposed mechanism can be tested in future experiments that monitor cancer tissues with live imaging to show the temporal co-evolution of fibroblasts and ECM network or in experiments that directly image deposited biopolymers in the culture experiments. The model would predict the co-evolution in the orientation of fibroblasts and corresponding ECM networks. On the other hand, if the accumulation of ECM materials or the interaction between fibroblasts and underlying ECM networks is significantly disrupted, the model would predict a diminished long-range alignment order of fibroblasts. Future efforts should focus on more detailed quantitative modeling of these mechanical interactions.

## Materials and Methods

### Modeling details of the kinetic Monte-Carlo simulations of elliptic particles

#### Motion of an elliptic particle along its major and minor axis

The length of the major and minor axis of an elliptic particle is 2*a* and 2*b*, respectively. Aspect ratio *r* = *a/b*. Along its major axis, a particle can move a distance *v · dt* in one simulation step. In our simulations, *dt* = 1 and *v* is a constant velocity along its major axis. At each time step, a particle can reverse its velocity with a probability *p* = 0.025. If possible, one particle is introduced into the system every simulation time step and one simulation time step corresponds to 0.05 hour in real time. This mimics the cell density increase observed in experiments. For simulations of elliptic particles, the self-propelled velocity is chosen as *v* = 0.1, 0.1667, 0.2333, 0.3 for different aspect ratios *r* = 3, 5, 7, 9, respectively.

Along its minor axis, a particle can randomly move with a step size *v_mi_* · *dt* · *v_mi_* is the velocity along its minor axis and is a random number between *−v*/*2r* and *v*/*2r*. In our simulations, varying the amplitude of *v_mi_* does not give rise to a perfect alignment of particles in a wide channel (Fig. S13). In addition, according to the movies in Ref. (10), an isolated fibroblast does not significantly trans-locate along its minor axis compared to its motion along its major axis.

#### Detecting overlaps between particles

In our kinetic Monte Carlo simulations, an elliptic particle is described by points on its boundary. If we define the angle *_pt_* as the angle between the semi-major axis and the line formed by a point on the boundary and the center of an ellipse, the angle difference Δ*θ_pt_* between adjacent points is the same.

In our simulations, the number of points for an elliptic particle with the aspect ratio *r* = 3, 5, 7, 9 are taken to equal 120, 240, 360, and 480, respectively. To detect whether there is any overlap between two particles, we simply calculate i) the distance *d_j_* between each point *j* on particle 1 and the center of particle 2; ii) the angle *θ_j_* between point *j* and the reference major axis of particle 2. If *d_j_* is smaller than the distance *d_j_′* of the point on particle 2 with the same angle *θ_j_*, the two particles should overlap with each other. We also do the same calculations for each point on particle 2 to further improve the accuracy of our detection. A detailed illustration of this procedure can be found in the Supplement Information Fig. S10.

We should note that our method is not exact. But it works well for the system we studied, which can ensure there are no particle configurations with any overlap between two particles in our simulations.

#### Rules of the interaction between channel boundaries and elliptic particles

After a proposed translational motion of a particle, if the particle moves out of the channel boundary and the particle is moving towards the boundary, first the particle will retreat back to its original location. Then a new orientation is proposed according to Eq. 1:

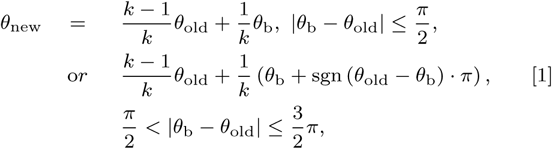

where θ_b_ = *π*/2 or 3*π*/2 for the left channel boundary or the right one, respectively; *k* is a random number evenly distributed between 2 and 10. In the simulation, we further check whether the particle with *θ*_new_ at its original location will overlap with its neighbors or not. If not, *θ*_new_ will be accepted and updated. If yes, all proposed motions fail and the particle simply keeps its original orientation and stays at its original location.

#### Rules of the reorientation after collisions between elliptic particles

When we examine a pair of particles with their newly proposed locations, if the two particles overlap with each other, first both particles will retreat back to their original locations. Then new orientations are proposed to reverse or rotate according to Eq. 2 with equal probability 0.5.

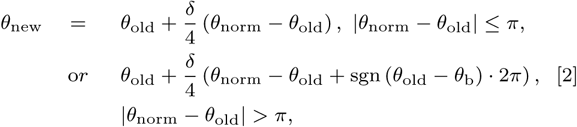

where *θ*_norm_ is the direction of the normal vector of the overlapped surface if the overlapped surface is in the head part of a particle; if the overlapped surface is in the tail part of a particle, *θ*_norm_ is the opposite direction of the normal vector of the overlapped surface. An illustration on how we decide *θ*_norm_ can be found in Fig. S11.

Furthermore, in Fig. S12 we show that varying the noise level *δ*/4 to δ, *δ*/2, or *δ*/8 cannot give rise to perfect alignment, where δ is an evenly distributed random number between 0 and 1.

#### Simulation procedures

i. propose new locations for all particles along their major and minor axes.
ii. for simulations carried out in a channel-like structure, we check whether each particle with the proposed location will run into boundaries or not. If yes, the particle retreats to its original location and rotates according to Eq. 1. If the particle will not overlap with its neighbors, the proposed orientation will be accepted and updated. If there is an overlap, the particle will simply keep its original orientation and location.
iii. we check whether particles will run into their neighbors. For particles with no neighbors around, proposed positions will be accepted and updated. For those with neighbors, it is then checked pair by pair in a sequential fashion whether particles will run into each other. If yes, both particles in a pair retreat to their original locations and then reverse or rotate according to Eq. 2 separately so that the two particles will be less likely overlapped further.
iv. with newly proposed orientations for the two particles in a pair, we then check that whether there is any further overlap between the first particle (*P*_1_) selected in the pair and all its neighbors (in their original positions and orientations). If there is no overlap, *P*_1_will get its new orientation accordingly. If there is an overlap, *P*_1_will keeps its original orientation and location. Then the second particle (*P*_2_) in the pair will be checked to see whether it will overlap with *P*_1_ (in the already-updated orientation) and all other neighbors. If there is no overlap, *P*_2_ will get its new orientation accordingly. If there is an overlap, *P*_2_ will keep its original orientation and location.

We perform procedure iii) and iv) within one simulation step in a sequential fashion for all pairs and we randomly permuted the sequence of particles after each simulation step.

#### Formulation of the orientation field and its effect on particles

To study the evolution of the “fiber” orientation, we first divide the 2-dimensional space into grids. The orientation field *θ* of each grid point will be affected by the orientations of particles *θ_n_* within a radius of *R_c_* surrounding that grid point in a time dependent manner. Specifically, the orientation field of each grid point will be updated according to Eq. 3.

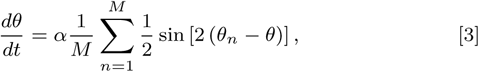

where *M* is the number of particles within a radius *R_c_* of each grid point. Fig. 5 illustrate the parameters involved. For small α (field update strength), the orientation field slowly responds to nearby fibroblasts and has a long memory of previous motion in its vicinity.

**Fig. 5.**
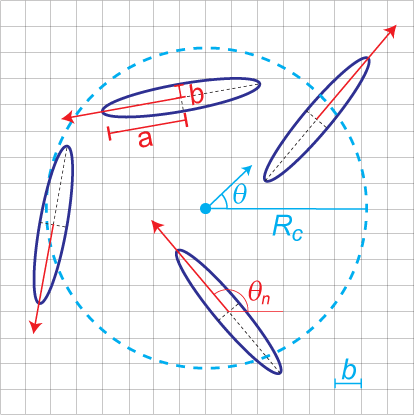
Illustration of our proposed mechanism. A cyan dot emphasizes a specific grid point on which we consider the orientation field. Elliptic particles falling within the dashed circle (radius, *R_c_*) will affect the orientation field at that grid point. The cyan arrow at the center of the circle indicates the current direction of that orientation field. Each particle’s current orientation is denoted by a red arrow from its center and it will bias the orientation field towards its direction; this is described quantitatively by Eq. (3).

For a given updated orientation field, a new orientation of each particle is proposed to equal *θ′_n_* where *θ′_n_* = *θ* if | *θ′_n_* − *θ*| ≤ π/2, or *θ* if |*θ′_n_* = *θ* − *π* if *π*/2 < *θ* − *θ′_n_* ≤ 3π/2, or *θ′_n_* = *θ* + π if π/2 < *θ′_n_* − *θ* ≤ 3π/2. *θ* is the orientation of its closest grid point of the orientation field.

The parameter *R_c_* gives the range in which the fiber orientation will be affected by the contraction of a fibroblast. There have been evidences showing that the contraction of fibroblasts can align neighboring fibers to the extent of a few times of the cell-body length (19, 33). Specifically, we chose *R_c_* as 3 times of the length of the semi-major axis of the fibroblast for the simulation in the channel-like confinement set-up. We also tested the effects of different *R_c_* on the nematic order of fibroblasts in a box with periodic boundary condition shown in Fig. S20.

### Sphero-cylinder cell model for fibroblasts

#### Cell movement

Here, fibroblasts are represented by a rectangle and two circles at two ends, i.e. a 2D sphero-cylinder. The particle width or the diameter of the circles at two ends is 2*b* and the particle length is *L_c_* = 2*rb*, where *r* is the aspect ratio. Each particle moves on a 2-D surface due to self-generated motility forces 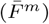 acting at the particle center in the direction of its current orientation *θ*. Further, particles periodically reverse (reversal period, *τ_R_*) their travel direction by flipping their current orientation by π (180-degrees).

#### Viscous drag

Fibroblast cells are surrounded by liquid medium and thus their motion is hindered by its viscous drag. We approximate the motility of cells as motion of particles in the over-damped limit (low Reynolds number regime) (34). We apply corresponding viscous drag forces 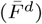 on particles, given by 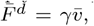, where 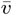 is the current velocity of the particle.

#### Cell-Cell collisions

We simulate excluded volume interactions during fibroblast movement through cell-cell collisions. Collisions in simulations are defined by the overlap of particle bodies during their movement. We resolve particle collisions by adding appropriate collision resolution forces 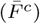 to each of the colliding particles at the point overlap in the direction of their collision normal 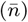.

#### Equation of motion

We solve the following equation of motion based on Newtonian dynamics under over-damped limit (linear - *γ* and angular - *ζ* drag coefficients) for each particle *i* in simulation subject to the forces acting on it and obtain its position 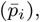, velocity 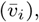 orientation (*θ_i_*) and angular velocity (*ω_i_*) at each step of the simulation.

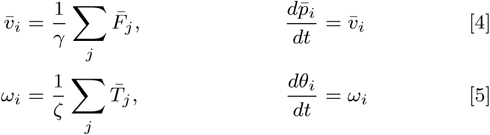

where summation is over various forces (*F_j_*) and torques (*T_j_*) acting on an particle. We use the Box2D (35) physics library to solve the equation of motion and resolve collisions in our simulation. All simulation parameters are given in Table S1.

#### Cell growth

Fibroblast cell density in experiments increased linearly with time when growing in channel structures (10) (except at initial and final stages of experiment). We replicate this linear cell density increase in simulation by adding new particles into simulation at a constant rate (*g*) equivalent to rate of cell density increase in experiments. We add new particles into the simulation by a selecting an existing particle randomly within the simulation region and initialize a new particle with same orientation as the selected particle. This process ensures that the added new particle does not disrupt the existing order in the system and imitates the cell division process in experiments (10) where daughter cells move with same orientation as the parent cell. The new particle position *p̄* is initialized randomly within a radius of *L_c_*/2 from the selected particle’s center. Additionally, cell density remained stationary in experiments after the density reached above 0.9 (10). We follow this process by stopping the addition of new particles into the simulation region when the system packing fraction (*η*) reaches close to 1.

## ACKNOWLEDGMENTS

This work was supported by the National Science Foundation Center for Theoretical Biological Physics (NSF PHY-1427654), and NSF DMS-1361411, Stand Up to Cancer, and The V Foundation. O.A.I. and R.B. are supported by NSF MCB-1411780. H.L. is also a Cancer Prevention and Research Institute of Texas (CPRIT) Scholar in Cancer Research of the State of Texas at Rice University. We thank the research associates Roger Wang and Anthony Rosario from P.L. group at the City of Hope for antibody staining and manuscript editing; the Ultima Intra-Vital system is supported by the Light Microscopy Core and Shared Resources at the City of Hope (Duarte, CA). We also thank Dr. Brian Armstrong (director of Light Microscopy Core at City of Hope) for editing part of the manuscript, and Erik Sahai, Paul Bates and Esther Wershof for insightful discussions.

